# MIDAS2: Metagenomic Intra-species Diversity Analysis System

**DOI:** 10.1101/2022.06.16.496510

**Authors:** Chunyu Zhao, Boris Dimitrov, Miriam Goldman, Stephen Nayfach, Katherine S. Pollard

## Abstract

**Summary:** The Metagenomic Intra-Species Diversity Analysis System (MIDAS) is a scalable metagenomic pipeline that identifies single nucleotide variants (SNVs) and gene copy number variants (CNVs) in microbial populations. Here, we present MIDAS2, which addresses the computational challenges presented by increasingly large reference genome databases, while adding functionality for building custom databases and leveraging paired-end reads to improve SNV accuracy. This fast and scalable reengineering of the MIDAS pipeline enables thousands of metagenomic samples to be efficiently genotyped.

**Availability and Implementation:** The source code is available at https://github.com/czbiohub/MIDAS2. The documentation is available at https://midas2.readthedocs.io/en/latest/.

**Supplementary Information:** Supplementary data are available at *Bioinformatics* online.

## 1 INTRODUCTIONW

Metagenotyping, the identification of intraspecific genetic variants in metagenomic data, is a powerful approach to characterizing population genetic diversity in microbiomes. Most pipelines identify variants based on alignment of reads to reference databases of microbial genomes and/or gene sequences (Supplementary Figure S1). While comprehensive reference databases can reveal strain-level relationships which would be otherwise overlooked (Beghini et al. 2021), alignment to large databases is computationally intensive. Furthermore, the divergence of reference genomes from strains in the metagenomic sample affects sensitivity and precision (Bush et al. 2020; Olm et al. 2021), and existing metagenotyping tools do not automatically adapt database files based on information in the metagenome. In this paper, we introduce MIDAS2 (Supplementary Figure S2), a major update to MIDAS (Nayfach et al. 2016) (Supplementary Table S1) that addresses these challenges through (1) a new database infrastructure geared to run on AWS Batch and S3 that achieves elastic scaling for constructing database files from large collections of genomes; and (2) a fast and scalable implementation of the SNV calling pipeline that enables metagenotyping in thousands of samples with improved accuracy achieved through utilization of paired-end reads and databases customized to the species present in the samples. As the only tool that integrates all steps of the metagenotyping process, from database customization to alignment and variant calling, MIDAS2 helps to promote reproducible research.

## 2 IMPLEMENTATION

### 2.1 MIDAS Reference Database Updates

We generated MIDAS Reference Databases (MIDAS DBs), comprised of species pangenomes, marker genes, and representative genomes, from two public microbial genome collections: UHGG v.1 (Almeida et al. 2021) (4,644 species / 286,997 genomes) and GTDB v202 (Parks et al. 2022) (47,893 species / 258,405 genomes). This is a significant increase in database content compared to MIDAS DB v1.2 (5,952 species / 31,007 genomes) and other tools (Supplementary Table S2). We implemented a new infrastructure that dramatically simplifies building a new MIDAS DB for other genome collections by using a table-of-contents file assigning genomes to species and denoting the representative genome for each species (Supplemental Fig S3). MIDAS DBs can be built locally, which enables customized selection of representative genomes, a key component of accurate SNV calling (Bush et al. 2020).

### 2.2. MIDAS2 Analysis Algorithm Updates

Metagenotyping SNVs across large numbers of samples is computationally intensive. First, alignment and pileup are applied to each species in each sample (single-sample step). Then these pileup results must be scanned for each genomic site to compute population SNVs (across-samples step). Metagenotyping methods such as MIDAS and metaSNV v2 parallelize these computations over species, capping the maximum number of processors (CPUs) at the number of species being genotyped. The SNV module of MIDAS2 achieves better CPU utilization by splitting genomic sites into multiple chunks per species. We execute parallelization over chunks in a way that does not destroy cache coherence to the point where computation stalls on input or output (I/O; Supplementary Notes).

## 3 RESULTS

### 3.1 Performance Improvements

We compared the running time and memory utilization of the single-sample and across-samples SNV modules of MIDAS and MIDAS2, using the same database (MIDAS DB v1.2) and 211 samples from an inflammatory bowel disease cohort (NCBI accession: PRJNA400072). The single-sample SNV module of MIDAS2 is slightly faster than MIDAS (Supplementary Figure S4), with database customization and Bowtie2 alignment taking up to 75% of run time (Supplementary Figure S5). The across-samples SNV module benefited more from parallelization, running 2.33 times faster in MIDAS2 with 48 CPUs (Figure 1A) and scaling linearly (Supplementary Figure S4).

**Figure 1.**
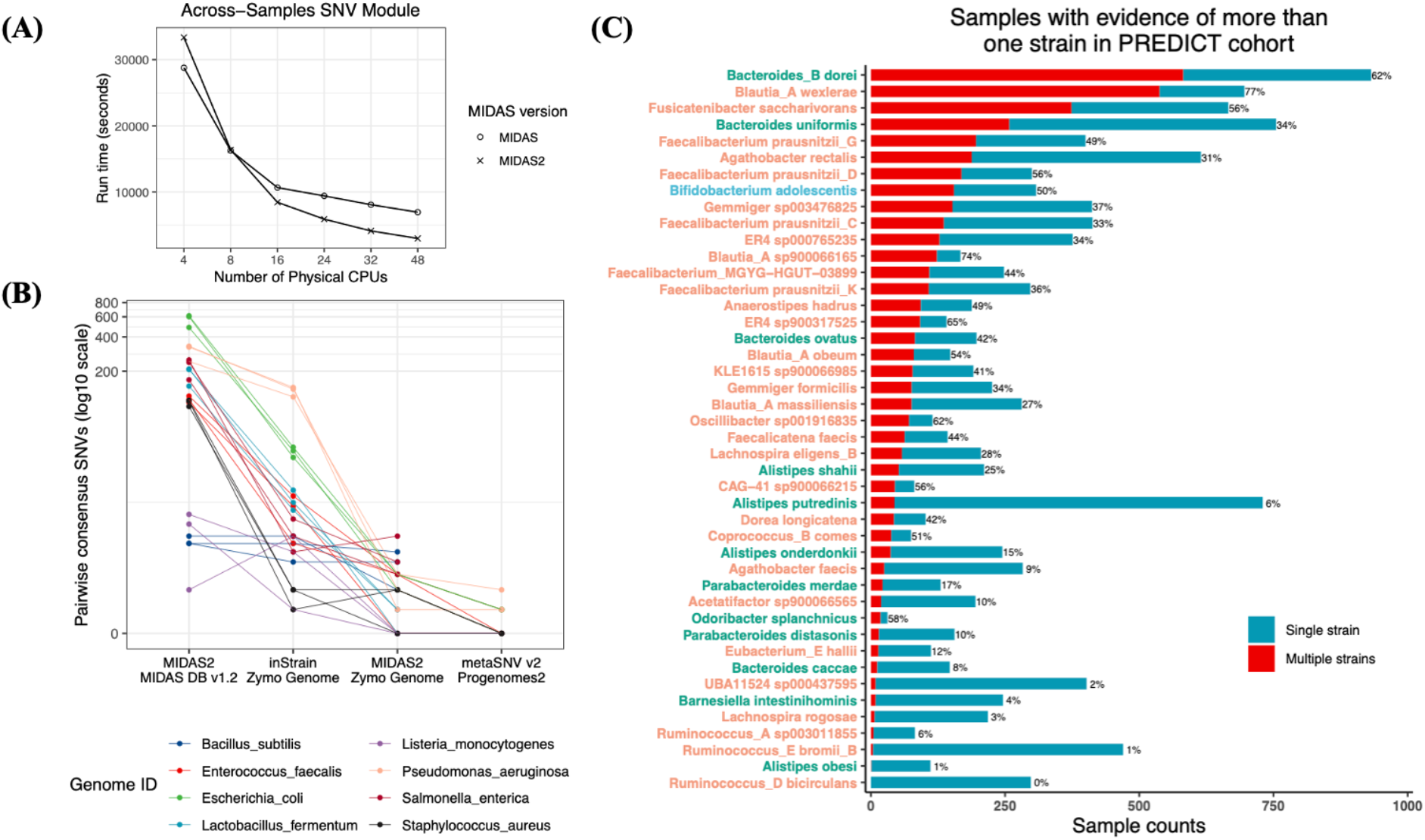
Speed, accuracy, and application of MIDAS2. **A:** The SNV module of MIDAS2 was re-engineered to parallelize within species, making it increasingly faster than MIDAS as more CPUs are deployed. This analysis was performed with 211 metagenomic samples (NCBI accession: PRJNA400072) and population SNVs. **B:** Benchmarking with three aliquots of a standardized microbial community shows that reference genome selection and post-alignment filters determine metagenotype accuracy. The aliquots are identical, so all SNVs are false positives. This analysis uses consensus SNVs. Population SNVs show the same pattern (Supplemental Figure S4). **C:** The distribution of samples with evidence of a strain mixture versus one dominant strain for 44 species metagenotyped by MIDAS2 in the PREDICT cohort (NCBI accession: PRJEB39223).

### 3.1 Reference Genomes and Post-alignment Filters Determine Metagenotype Accuracy

MIDAS2, inStrain and metaSNV v2 were applied to three aliquots of a standardized bacterial community (Olm et al. 2021), and SNVs were compared between aliquots which should have identical metagenotypes (Supplementary Note). metaSNV v2 has the fewest false positives by only using uniquely aligned reads, but it only genotyped five of the eight species in the community (Supplementary Table S5). InStrain and MIDAS2 correctly detected all eight species. When both are run with a genome database containing only the reference genomes of the strains in the community, MIDAS2 has fewer false positives (Figure 1B). However, the false positive rate of MIDAS2 is higher when using the MIDAS DB v1.2, in which these species’ reference genomes are diverged from the sample. Thus, high-quality reference genomes and post-alignment filters that balance false positives against false negatives are crucial for metagenotyping.

### 3.3 Scalability of MIDAS2 versus metaSNVv2

Since metaSNV v2 was previously shown to be efficient enough to metagenotype thousands of samples (Van Rossum et al. 2021), we assessed the scalability of MIDAS2 compared to metaSNV v2 on 1,097 samples from the PREDICT study (NCBI accession: PRJEB39223), using MIDAS DB UHGG with both tools (Supplementary Note). Despite the same species selection criteria, MIDAS2 metagenotyped many more species (44 versus 14 for metaSNV v2) (Supplementary Note). MIDAS2 used more memory (21.21 GB versus 4 GB peak RAM utilization) and ran slightly longer (average 106 versus 84 minutes per species) to achieve this. We conclude that MIDAS2 can metagenotype thousands of samples with reasonable computational costs, providing a more sensitive alternative to metaSNV v2.

### 3.4 MIDAS2 Detects Metagenomes with Multiple Strains

For each of the 44 species from PREDICT with MIDAS2 metagenotypes, we quantified evidence of a single dominant strain versus mixtures of multiple strains with an existing method (Nayfach and Pollard 2015; Garud et al. 2019; Nayfach et al. 2016). While most species showed evidence of distinct lineages across samples (Supplementary Figure S8), single samples often had a single dominant strain (Figure 1C). However, samples with strain mixtures were common for several species, including *Bacteroides_B dorei* (62%) and *Faecalibacterium prausnitzii_G* (49%) (Supplementary Figures S9 and S10).

## ACKNOWLEDGMENTS

This work was funded by the Chan Zuckerberg Biohub, Gladstone Institutes, and NSF grant #1563159. Conflicts of Interest: none declared.

## Supplementary Note

### MIDAS2: Strain-level Reads-to-Table Metagenotyping Pipeline

MIDAS2 performs strain-level metagenomics analysis through a SNV module and a CNV module. Each module includes two sequential steps: single-sample analysis and across-samples integration of single-sample results to identify population variants (Supplementary Figure S2). The SNV module is based on alignment of metagenomic reads to one representative genome per species (rep-genomes), and the CNV module is based on alignment of reads to pan-genomes; both use an estimate of which species are present in the sample to customize the sample-specific reference database.

Given a collection of shotgun metagenomics samples, both the SNV and CNV module start with identifying the list of species that are sufficiently abundant for metagenotyping, based on profiling 15 universal, single copy genes (SCGs). By default, all species with median SCG coverage > 2X are metagenotyped. If the purpose is to genotype low abundance species, users are responsible for adjusting the cutoff (e.g., median SCGs coverage > 0). Species to be metagenotyped are then used to construct a sample-specific rep-genome database that only contains the representative genomes for these species (SNV module) and/or a pan-genome database that only contains the pan-genomes for these species (CNV module). This customization of a pre-computed MIDAS DB to the sample improves metagenotype accuracy by reducing “cross-mapping”, whereby metagenomic sequencing reads align erroneously to conserved regions in the genomes of closely related species.

The SNV module starts with single-sample pileup, by aligning reads to a sample-specific rep-genome database. The alignment results for all the species in the rep-genome database are reported, such as *fraction_covered mean_coverage*, and *mapped_reads*. MIDAS2 purposely holds off on any filtering or species selection with the single-sample pileup results until across-samples SNV analyses are performed. Upon the completion of the single-sample SNV analyses for all the samples, MIDAS2 will compute the population SNVs across all the samples (see Population SNV Computation section).

It is worth noting that the MIDAS2 single-sample SNV module is not designed for consensus SNV calling, and by design MIDAS2 does not apply any species or variants calling filters at the single-sample level. However, in recognition of the need for single-sample consensus SNV calling, we added an *-- advanced* mode to the single-sample SNV analysis in MIDAS2 to report per species major allele and minor allele for all the genomic sites covered by at least two reads, upon which any variant calling filter would be applied by the users. We highly recommend users set the parameter *--ignore_ambiguous* to ignore ambiguous alleles, where genomic sites recruit tied read counts.

The CNV module starts with single-sample alignment of reads to the sample-specific pan-genome database. As in the SNV module, this only includes the pan-genomes of species being genotyped (i.e., sufficiently abundant species based on median SCGs coverage). The alignment summaries for all the species in the customized pan-genome database are reported (e.g. *mean_coverage, mapped_reads*), and these are used to determine the scope of across-samples CNV analysis. For example, only species that pass vertical pan-gene read depth > 1 (by default) are included in the across-samples CNV computation. For each gene in each species, coverage is normalized by the mean coverage of all the SCGs to estimate copy number per cell (Nayfach and Pollard 2016). A brief comparison of the CNV module with other methods can be found in Supplementary Table S3.

### Features Distinguishing Different Reference-based Metagenotyping Pipelines

All read mapping based metagenotyping pipelines share an underlying structure, yet they differ from one another in the availability of genome databases, the customization of reference genomes to the species present in the sample, read aligners, pileup tools, post-alignment filters, and the contents of input and output files (Supplementary Figure S1).

#### Database

Whole-genomes are used as the reference for most pipelines (MIDAS2, metaSNV v2 (Van Rossum et al. 2021), InStrain (Olm et al. 2021)), while species-specific marker genes are used by StrainPhlan (Truong et al. 2017). MIDAS2 is the only pipeline providing precomputed reference genome databases: namely MIDAS DB UHGG and MIDAS DB GTDB. MIDAS2 builds a database of representative genomes (rep-genomes) customized to only include species that are sufficiently abundant in the sample, which is determined via coverage of 15 SCGs. In contrast, metaSNV v2 and inStrain leave database selection and customization to their users.

#### Alignment and pileup

MIDAS2 and inStrain provide post-alignment filters based on the read alignment quality (e.g., alignment sequence similarity and uniqueness), while metaSNV v2 leaves this step to the user. MIDAS2 performs read pileup directly from counting reads covering each nucleotide (*count_coverage*), while metaSNV v2 and inStrain both use samtools *mpileup* (Danecek et al. 2021) or pysam (Andreas Heger et. al. 2021) for the pileup task. See Supplementary Table S12 for the resulting differences in variant calling results. One possible reason for differences in metagenotypes is that samtools does not use soft-clipped reads. This may be desirable for certains applications. Yet since all microbiome pipelines apply post-alignment filtering to the raw alignment file before the pileup and variant calls, an unfiltered read pileup seems to be appropriate, so this is what we implemented in MIDAS2.

#### Genotypes

For single-sample SNV calling, inStrain only reports non-reference variants (when the major allele is different from the reference allele), which can cause missing allele information for downstream across-samples population SNV calling. On the other hand, metaSNV v2 does not report any single-sample pileup results, and only reports the across-samples non-reference variants. Adding samples to an existing analysis, metaSNV v2 therefore has to be run again from the beginning. MIDAS2 reports single-sample pileup results for the all genomic sites covered by at least 2 reads. Therefore, MIDAS2 can avoid repeating single-sample pileup when more samples are added to the analysis. MIDAS2 reports population SNVs of all genomic sites meeting the user-defined types (e.g, *snp_type*).

In summary, MIDAS2 is the only reads-to-table metagenotyping pipeline. By integrating and automating all steps of the metagenotyping process, MIDAS2 helps to promote reproducible research.

### Population SNV Computation

Population SNV analysis (across-samples step) restricts attention to “sufficiently well” covered species in “sufficiently many” samples. To be specific, a given *<species, sample>* pair will only be kept if it has more than 40% horizontal genome coverage (*fraction_covered*) and 5X vertical genome coverage (*mean_coverage*). Furthermore, only “sufficiently prevalent” species with “sufficiently many’’ samples (by default *sample_counts* > 2) are eligible for across-samples population SNV analyses. Therefore, different species may have different lists of relevant samples. For each genomic site, a sample is “relevant” if the corresponding site depth falls between the user-defined range, otherwise it is ignored for the pooled-SNV compute. Therefore, different genomic sites from the same species may have different panels of “relevant samples”. And genomic site prevalence can be computed as the ratio of the number of relevant samples for the given site over the total number of relevant samples for the given species.

There are three main steps to compute and report population SNV in MIDAS2. First, for each site, MIDAS2 determines the set of alleles present across all relevant samples. Specifically, for each allele (A, C, G, T), *merge_snps* subcommand (1) tallies the sample counts (*sc*) of relevant samples containing corresponding allele, and (2) sums up the read counts (*rc*) of the corresponding allele across all the relevant samples. Second, population major and minor alleles for a single site can be computed based on the accumulated read counts or sample counts across all relevant samples. The population major allele refers to the most abundant/prevalent allele, and the population minor allele refers to the second most prevalent/abundant allele. Third, MIDAS2 collects and reports the sample-by-site matrix of the corresponding (1) site depth and (2) allele frequency of the population minor allele for all the relevant samples

### Chunkified Pileup Implementation

MIDAS parallelized pileup work by species, and because some species required more work than others, this resulted in a few CPU cores working longer while the rest sat idle waiting. So MIDAS2 subdivided the work for those large species into smaller units that we called chunks, by splitting the species’ genomes into segments. We size the chunks so they take reasonable time (default *chunk_size* = 1000000). If chunks are too big, we will have CPU cores sitting idle while others finish running. If chunks are too small, the per-chunk setup costs will be high. The chunkified pileup algorithm can sweep through a large data set coherently, ensuring data locality so the active chunks fit into available filesystem cache, and it therefore achieves better utilization.

To compute the per-chunk population SNV, all the pileup results of corresponding sites across all the samples need to be read into memory. Therefore at any given moment, up to total CPU cores * total number of sites per chunk * total number of the relevant samples pileup results will be read into RAM. Therefore, for a highly prevalent species, MIDAS2 dynamically (--*robust_chunk*) adjusts to a smaller chunk size. There is a trade-off between the chunk size, the running time and the memory usage. Users can customize the chunk size according to their computing environment. For example, constant RAM usage can be achieved despite an increasing number of samples with dynamic chunk sizes. When all chunks from the same species finish processing, chunk-level pileup results will be merged into species-level pileup results. Below is the pseudocode for the chunkified population SNV implementation.

~~~
def process(species_id, chunk_id):
   if chunk_id == -1:
     collect_chunks()
   chunk_worker()
def chunk_worker():
   for sample in list_of_relevant_samples:
       accumulate()
   call_population_snps()
   write_population_snps()
multiprocessing_map(process, list_of_species, num_cores)
~~~

This implementation makes reference-based population SNV analysis across thousands of samples possible.

### Consensus and Population SNVs for a Standardized Microbial Community Reveal that Reference Genome Selection and Post-alignment Filters Determine Metagenotyping Accuracy

We compared the consensus sequences reported by three metagenotyping pipelines using three metagenomes from aliquots of the same ZymoBIOMICS Microbial Community Standard product (catalog no. D6300, BioProject PRJNA648136). No genetic differences were expected between these samples. For each species, SNVs were called by comparing all pairs of metagenomes; and we computed the pairwise consensus SNVs and population SNVs to assess the false positive rates of each method as in (Olm et al. 2021).

#### Analysis

Three metagenotyping pipelines that use alignment to reference databases were compared: MIDAS2, inStrain, and metaSNV v2. We ran inStrain with the Zymo reference genomes as in (Olm et al. 2021), metaSNV v2 with proGenomes 2 as in (Mende et al. 2020), and MIDAS2 with MIDAS DB v1.2 for the purpose of reproducibility with results in (Olm et al. 2021). For the post-alignment filters, we used the default or recommended filter parameters by each tool (Supplementary Table S4). To evaluate the role of a well-matched reference database, we also ran MIDAS2 with the Zymo reference genomes, while keeping everything else unchanged. This allowed for a direct comparison of the post-alignment filters deployed in inStrain and MIDAS2, as well as a direct assessment of the reference database between MIDAS DB v1.2 and Zymo reference genomes. All the tools report horizontal genome coverage, upon which species presence is called. The same set of variant calling filters as inStrain were applied to all the tools. Last, pairwise consensus-based SNVs (consensus SNVs) and population SNVs (Olm et al. 2021) were computed. Simply put, consensus SNVs refer to two major alleles that are different between a pair of samples, whereas population SNVs refer to sites that differ (neither major or minor allele match) between a pair of samples.

#### Database

For the eight bacterial Zymo reference genomes, we found the corresponding species in MIDAS DB v1.2 and proGenomes2 based on the highest Average Nucleotide Identity (ANI) (Jain et al. 2018). It is worth mentioning that for Lactobacillus fermentum and Salmonella enterica, there are two pairs of highly closely related strains in the proGenomes2: 98.8% and 98.5% (Supplementary Table S6). We also reported the genome quality of all the representative genomes in all three databases using checkM (Parks et al. 2015) (Supplementary Table S7).

#### Results

The three aliquot samples originated from the same microbial population, therefore any observed SNVs could be a product of (1) read misalignment: some regions of the reference genome erroneously recruits reads originating from other genome regions or even other species; (2) failure of the post-alignment filters to recognize the misalignments as such; or (3) microdiversity created in the lab.

MIDAS2 reported more accurate consensus sequences compared to inStrain when both tools were run using the Zymo reference genomes matched to the strains in the metagenomes (Figure 1B). However, the accuracy of MIDAS2 was much inferior when the Zymo reference database was replaced by MIDAS DB v1.2, in which the representative genomes are less similar to the community and also suffer from fragmented assemblies. Population SVNs followed the same pattern (Supplementary Figure S6). This shows that the reference genomes are the major determinant of accuracy for consensus SNVs, explaining a previous report of poor performance for MIDAS (Olm et al. 2021). As suggested by Olm et al., a good representative genome has the following characteristics: high quality contiguous sequence and a high degree of shared gene content with the taxa it is meant to represent.

On the other hand, with its stringent post-alignment filters (e.g., recruiting only uniquely aligned reads), metaSNV v2 only reported metagenotypes for five of the eight species in the mock community. Three species did not pass its default 40% horizontal coverage threshold (Supplementary Table S8). A few reasons can explain this failure of metaSNV v2. First, there is not an appropriate representative genome of the species *Bacillus subtilis* in the proGenomes 2; the closest genome has average nucleotide identity (ANI) = 92.54% (txid 224308). Second, there are two very closely related genomes in the proGenome 2 database for the other two species: *Lactobacillus fermentum* (ANI 98.83%) and *Salmonella enterica* (ANI 98.53%), which led to extremely low numbers of uniquely alignable reads for these two species. Since metaSNV v2 requires uniquely mapped reads, these two species have low horizontal genome coverage (0.04% and 30% of the whole genome, respectively) and hence many fewer sites metagenotyped. Third, metaSNV v2 only reports non-reference variants in the output. Therefore significantly fewer genomic sites were metagenotyped and further compared between a pair of samples by metaSNV v2. As a result, metaSNV v2 ran faster than the other two tools and reported fewer false positive SNVs (Supplementary Table S5). For the 1097 samples from the PREDICT study (PRJEB39223), the analysis-ready BAM file of MIDAS2 is 1.53 times larger than metaSNV v2, again due to the different post-alignment filters. Thus, metaSNV v2 generates few false positive SNVs and runs efficiently, but it has lower sensitivity than MIDAS2 and inStrain.

To summarize, we demonstrated building a high quality genome database and utilizing post-alignment filters that balance false positive versus false negative SNVs determine metagenotype accuracy.

### MIDAS2 Reference Database Target Layout and Construction

MIDAS Reference Database (MIDAS DB) refers to a set of custom files needed to run MIDAS2. There are three components in a MIDAS DB: rep-genome database, SCG marker database, and pan-genome database (Supplemental Figure S3). In each part, data is organized by species. For a given species, the rep-genome database contains all representative genomes of that species annotated using Prokka (Seemann 2014), and the SCG marker database is built on identified homologs of 15 universal SCGs from the representative genomes. The pan-genome database refers to the set of non-redundant genes within all genomes from that species (Commichaux et al. 2021), clustered at 99% sequence identity using vsearch (Rognes et al. 2016). The original release of MIDAS provided a default bacterial reference database (MIDAS DB v1.2), constructed from a collection of 5952 bacterial species clusters representing 31,007 high-quality bacterial genomes. With the rapid growth of the number of sequenced microbial genomes, particularly with the addition of metagenome-assembled genomes (MAGs) from diverse habitats, it is necessary to update the MIDAS DB.

For MIDAS2, we took advantage of several published collections of prokaryotic genome collections (UHGG (Almeida et al. 2021), GTDB (Parks et al. 2022)) in which genomes were already clustered into species groups. We developed a new database infrastructure that is geared to run on AWS Batch and S3, achieving elastic scaling for database construction for each of these large genome collections. Specifically, MIDAS DB construction can be executed in AWS using hundreds of instances, depositing built products in S3. For example, the species pan-genome for all 47,894 species of GTDB r202 was built in roughly one week at a cost of $80K, using 100 r5d.24xlarge instances. The new database infrastructure reads in a table-of-contents (TOC) file containing genome-to-species assignments for all the genomes and a choice of representative genome for each species cluster. Six-digit numeric species ids are randomly assigned and stored in the corresponding metadata file (*metadata*.*tsv*). MIDAS2 users can also build a new MIDAS DB locally for a small collection of representative genomes of interest.

### UHGG Separable/Inseparable Species

Many species have a closely related species with pairwise average nucleotide identity (ANI) near the species boundary (e.g., 95% ANI). For example, in the UHGG v1 database (Almeida et al. 2021), there are 981 species with closest pairwise ANI higher than 92%. We explored how effective SCGs are in separating the species in the UHGG database. For a given SCG in a given species, we compute all 31-mers and compare them to 31-mers present in all species. We define the marker as separable if it has >100 unique 31-mers. We then define a species as separable if more than 50% of its 15 SCGs are separable. We found that there are 3,956 separable species and 689 inseparable species (Supplementary Tables S13 and S14). The majority (76%) of inseparable species are from the *Collinsella* genus. The representative genomes of the 3,956 separable species were used as the reference database for the PREDICT analysis in this study.

### Bioinformatics Processing of Publicly Downloaded Samples

Metagenomic sequencing reads from an inflammatory bowel disease cohort (NCBI accession: PRJNA400072) were pre-processed using Sunbeam (Clarke et al. 2019), and 211 samples with more than 5 million reads were used for benchmarking. The benchmark work was done on a m5.16xlarge or m5.24xlarge EC2 instance, depending on the CPU counts needed.

Metagenomic sequencing reads from the PREDICT cohort (NCBI accession: PRJEB39223) were pre-processed using Sunbeam (Clarke et al. 2019), and 1,097 samples with more than 5 million reads were used in our analysis. The benchmark of MIDAS2 and metaSNV v2 was done on a r5-24xlarge instance.

The bioinformatics details of the ZymoBIOMICS benchmarking experiment are shown in Supplementary Table S4. The parameters used in the PREDICT analysis are shown in Supplementary Table S10.

### Parameters for paired-end read alignment in MIDAS2

During Bowtie2 read alignment, the default maximum fragment length is 500 base pairs (bp). With this setting, any read pairs sequenced from DNA fragments longer than 500 bp are labeled as not properly aligned (*is_proper_pair* = FALSE), and the reported template length may also be incorrect (equal to zero) (Supplementary Figure S12). Therefore, to properly use the paired-end option of the single-sample SNV module, it is crucial to set the *-X* (maximum fragment length) to an appropriate value for the metagenomes being analyzed (default =5,000 bp), even if longer maximum fragment length will make the alignment slower.

### Quasiphasable Species Model

We applied the model from Garud, Good et al. (Garud et al. 2019) to the PREDICT study to determine whether each species in each metagenomic sample was quasi-phasable (QP) or not (i.e., one dominant strain versus colonization by multiple bacterial lineages). The model uses synonymous sites in genes of the core genome of a given species. If the fraction of these sites with intermediate allele frequencies is high, this is taken as evidence for a strain mixture.

#### Bioinformatics

We use similar sample and site filters for population SNVs as in (Garud et al. 2019). Specifically, we filter the per species *snps_info*.*tsv* and *snps_freqs*.*tsv* files as follows:

- Minimal per sample median site depth of bi-allelic SNVs from protein coding sequences 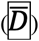 is 20
- Only include 4-fold degenerate synonymous sites
- Sample site depths must be between. 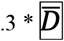 and 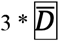
- Minimal site depth is 10
- Minimal site prevalence is 5

This produces a filtered SNVs allele frequency file.

#### QP calculation

For each species, we estimated whether each sample is QP as follows:

1. For each non-intermediate site, we defined population major allele direction 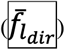 as 1 if the majority of the allele frequencies are higher than 0.8 and 0 if not.

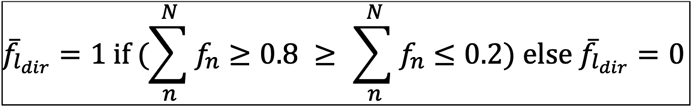

where 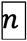 is the sample index, 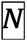 is the number of relevant samples for the given site.
2. For each sample 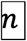, we computed the dominant haplotype of each non-intermediate site 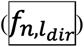 as 1 if the corresponding allele frequency is higher than 0.8 and 0 if not

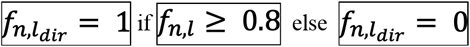

where 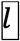 is the site index.
3. For each non-intermediate site, the population major allele frequency 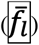 is computed as:

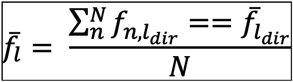
4. For each sample, we estimated 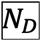 as the average genetic distance between sample *n* and the alleles present in the remainder of the samples 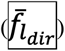 as:

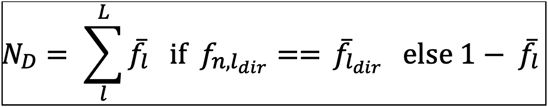

where, 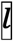 is site index, is 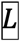 total number of non-intermediate sites; 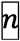 is the sample index.
5. For each sample, we computed the number of intermediate alleles over all sites 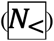 :

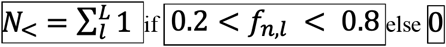
6. For each sample, if 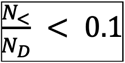 then we say the sample is QP, because there is evidence of one dominant strain with sufficiently high coverage and sufficiently low rates of intermediate alleles.

### PCoA and Manhattan distance

Manhattan distance was calculated based on the filtered SNV site-by-sample allele frequency matrix to evaluate the dissimilarity between samples. Principal Coordinate Analysis (PcoA) was calculated based on the Manhattan distance matrix using the ape package (Paradis et al. 2004).

